# Characterisation of AmpC Hyper-Producing *Escherichia coli* from Humans and Dairy Farms Collected in Parallel in the Same Geographical Region

**DOI:** 10.1101/784694

**Authors:** Maryam Alzayn, Jacqueline Findlay, Hannah Schubert, Oliver Mounsey, Virginia C. Gould, Kate J. Heesom, Katy M. Turner, David C. Barrett, Kristen K. Reyher, Matthew B. Avison

## Abstract

**Objectives:** To characterise putative AmpC hyper-producing 3^rd^ generation cephalosporin-resistant *E. coli* from dairy farms and their phylogenetic relationships as well as to identify risk factors for their presence; to assess evidence for their zoonotic transmission into the local human population

**Methods:** Proteomics was used to explain differences in antimicrobial susceptibility. Whole genome sequencing allowed phylogenetic analysis. Multilevel, multivariable logistic regression modelling was used to identify risk factors.

**Results:** Increased use of amoxicillin-clavulanate was associated with an increased risk of finding AmpC hyper-producers on farms. Expansion of cephalosporin resistance in AmpC hyper-producers was seen in farm isolates with *marR* mutations (conferring cefoperazone resistance) or when AmpC was mutated (conferring 4^th^ generation cephalosporin and cefoperazone resistance). Phylogenetic analysis confirmed the dominance of ST88 amongst farm AmpC hyper-producers but there was no evidence for acquisition of farm isolates by members of the local human population.

**Conclusions:** In this two-year surveillance study of 53 dairy farms, AmpC hyper-production was the cause of cefotaxime resistance in 46.2% of *E. coli*. There was evidence of recent farm-to-farm transmission and of adaptive mutations to expand resistance. Whilst there was no evidence of isolates entering the local human population, efforts to reduce 3^rd^ generation cephalosporin resistance on dairy farms must address the high prevalence of AmpC hyper-producers. The finding that amoxicillin-clavulanate use was associated with increased risk of finding AmpC hyper-producers is important because this is not currently categorised as a highest-priority critically important antimicrobial and so is not currently targeted for specific usage restrictions in the UK.

## Introduction

*Escherichia coli* typically produce a class 1 cephalosporinase, encoded by the *ampC* gene, which is chromosomally located. Expression of *ampC* in wild-type cells is low and not enough to confer clinically relevant resistance to β-lactam antibiotics. Many mutations, insertions and gene duplication events have been shown to cause *ampC* hyper-expression, and this leads to varying spectra of β-lactam resistance, dependent on the actual amount of AmpC produced.^1^ AmpC hyper-production was first seen in *E. coli* from human clinical samples in 1979,^2^ and for a period before the emergence of plasmid-mediated extended spectrum β-lactamases, AmpC hyper-production was a dominant mechanism of 3^rd^ generation cephalosporin (3GC) resistance in *E. coli* from humans.^1^ This is no longer the case, however. For example, in a recent survey of cefotaxime resistant (CTX-R) *E. coli* from urine collected from people living in South West England, only 24/626 isolates (3.8%) were presumed to be AmpC hyper-producers because of their lack of horizontally acquired β-lactamase genes; whole genome sequencing (WGS) confirmed that 13/13 sequenced isolates had *ampC* promoter mutations typical of AmpC hyper-producers.^3^

AmpC is typical of class 1 enzymes in that it does not confer resistance to the 4^th^ generation cephalosporins (4GC).^1^ However, *ampC* structural variants of *E. coli*, expanding AmpC activity to include, for example, cefepime, have been identified from humans ^4–7^ and cattle;^8^ these are dominated by isolates from the relatively less pathogenic phylogroup A, and particularly ST88,^6, 8^ probably because expanded-spectrum activity evolves from existing AmpC hyper-producers, of which ST88 isolates are particularly common.^9^

We recently conducted a survey of 4594 samples collected from faecally contaminated sites on 53 dairy farms in South West England. We identified 384 samples, collected across 47 farms, that were positive for the detectable growth of CTX-R *E. coli* isolates.^10^ In an recent paper, we reported that 566/1226 of these CTX-R *E. coli* isolates (from 186 samples from 38 farms) were PCR-negative for mobile cephalosporinases and so were presumed to be chromosomal AmpC hyper-producers.^11^ If this presumption was correct, AmpC hyper-production was the mechanism of resistance in 46.2% of CTX-R *E. coli* from dairy cattle in this region of the UK. This figure is comparable with the 42.9% presumed AmpC hyper-producers seen in CTX-R *E. coli* from dairy cattle in a recent nationwide Dutch study ^12^ and contrasts with the 3.8% of AmpC hyper-producers seen in CTX-R isolates in our recent study of human urinary *E. coli*.^3^

One aim of the work reported here was to characterise putative AmpC hyper-producing *E. coli* from our recent survey of dairy farms ^10, 11^ and to identify risk factors for the presence of AmpC hyper-producers on these farms. Another aim was to investigate potential zoonotic transmission of AmpC hyper-producers by using WGS-based phylogenetic analysis to compare isolates from farms located within an approximately 50 × 50 km sub-region of the study with human urinary *E. coli* collected in parallel from this same sub-region.^3^

## Materials and Methods

### Bacterial isolates, identification and susceptibility testing

Isolates used in this study came from dairy farms located within a sub-region of the wider study area.^10,11^ This region was chosen because it also included the locations of 146 GP practices that submitted urine samples for processing at the Severn Pathology laboratory, as described in a recently published survey of human urinary *E. coli*;^3^ this laboratory was also the source of the human urinary isolates used in the present study. Disc susceptibility testing and microtiter MIC assays were performed and interpreted according to CLSI guidelines.^13–15^

### Fluorescent Hoescht (H) 33342 dye accumulation assay

Envelope permeability in living bacteria was tested using a standard dye accumulation assay protocol ^16^ where the dye only fluoresces if it crosses the entire envelope and interacts with DNA. Overnight cultures in Cation Adjusted Muller Hinton Broth (CA-MHB) at 37°C were used to prepare CA-MHB subcultures, which were incubated at 37°C until a 0.6-0.8 OD_600_ was reached. Cells were pelleted by centrifugation (4000 rpm, 10 min; ALC, PK121R) and resuspended in 1 mL of phosphate-buffered saline. The optical densities of all suspensions were adjusted to 0.1 OD_600_. Aliquots of 180 μL of cell suspension were transferred to a black flat-bottomed 96-well plate (Greiner Bio-one, Stonehouse, UK). Eight technical replicates for each strain tested were in each column of the plate. The plate was transferred to a POLARstar spectrophotometer (BMG Labtech) and incubated at 37°C. Hoescht dye (H33342, 25 μM in water) was added to bacterial suspension of the plate using the plate-reader’s auto-injector to give a final concentration of 2.5 μM per well. Excitation and emission filters were set at 355 nm and 460 nm respectively. Readings were taken in intervals (cycles) separated by 150 seconds (s). Thirty-one cycles were run in total. A gain multiplier of 1300 was used. Results were expressed as absolute values of fluorescence versus time.

### Proteomics

1 mL of an overnight CA-MHB culture were transferred to 50 mL CA-MHB and cells were grown at 37°C to 0.6-0.8 OD_600_. Cells were pelleted by centrifugation (10 min, 4,000×g, 4°C) and resuspended in 35 mL of 30 mM Tris-HCl, pH 8 and broken by sonication using a cycle of 1 s on, 0.5 s off for 3 min at amplitude of 63% using a Sonics Vibracell VC-505TM (Sonics and Materials Inc., Newton, Connecticut, USA). The sonicated samples were centrifuged at 8,000 rpm (Sorval RC5B PLUS using an SS-34 rotor) for 15 min at 4°C to pellet intact cells and large cell debris. Protein concentrations in all supernatants were quantified using Biorad Protein Assay Dye Reagent Concentrate according to manufacturer’s instructions. Proteins (1 μg/lane) were separated by SDS-PAGE using 11% acrylamide, 0.5% bis-acrylamide (Biorad) gels and a Biorad Min-Protein Tetracell chamber model 3000X1. Gels were resolved at 200 V until the dye front had moved approximately 1 cm into the separating gel. Proteins in all gels were stained with Instant Blue (Expedeon) for 5 min and de-stained in water. LC-MS/MS data was collected as previously described.^17^ The raw data files were processed and quantified using Proteome Discoverer software v1.4 (Thermo Scientific) and searched against bacterial genome and horizontally acquired resistance genes using as described previously.^18^

### Whole genome sequencing and analyses

WGS was performed by MicrobesNG (https://microbesng.uk/) on a HiSeq 2500 instrument (Illumina, San Diego, CA, USA) using 2×250 bp paired end reads. Reads were trimmed using Trimmomatic^19^ and assembled into contigs using SPAdes 3.13.0^20^ (http://cab.spbu.ru/software/spades/). Resistance genes, plasmid replicon types and sequence types (according to the Achtman scheme^21^) were assigned using the ResFinder,^22^ PlasmidFinder,^23^ and MLST 2.0 on the Center for Genomic Epidemiology (http://www.genomicepidemiology.org/) platform. Contigs were annotated using Prokka 1.2.^24^

### Phylogenetic analysis

Sequence alignment and phylogenetic analysis was carried out on the Bioconda software package^25^ on the Cloud Infrastructure for Microbial Bioinformatics (CLIMB).^26^ Sequences were first aligned to a closed read reference sequence and analysed for SNP differences, whilst omitting insertion and deletion elements, using the ‘Snippy’ alignment program. Alignment was then focused on regions of the genome found across all isolates, using the Snippy-core program, thus eliminating the complicating factors of insertions and deletions.^27^ Aligned sequences were then used to construct a maximum likelihood phylogenetic tree using RAxML, utilising the GTRCAT model of rate heterogeneity and the software’s autoMR and rapid bootstrap to find the best-scoring maximum likelihood tree and including tree branch lengths, defined as the number of base substitutions per site compared.^28, 29^ Finally, phylogenetic trees were illustrated using the web-based Microreact program.^30^

### Risk factor analysis

Multivariable, multilevel logistic regression analysis was performed to identify risk factors for the presence of AmpC hyper-producers in samples collected from farms. Positivity for AmpC hyper-producing *E. coli* in a sample was defined by the growth of *E. coli* on cefotaxime agar which were PCR-negative for known horizontally-acquired cefotaxime resistance genes.^11^ The risk factor analysis methodology used has been described previously, including the use of a novel method using a logistic link function to account for measurement error.^10^

## Results and Discussion

### Confirmation of AmpC hyper-production and identification of porin loss and *marR* mutations in *E. coli* from dairy farms

In order to further investigate putative AmpC hyper-producing *E. coli* isolates from dairy farms identified in our recent surveillance study,^10,11^ antibiograms were determined for one putative AmpC hyper-producing isolate from each of 5 randomly selected farms. All isolates presented a typical AmpC-hyper-producing phenotype: resistance to ampicillin and cefalexin, and non-susceptibility to cefotaxime and ceftazidime. The isolate from Farm 1 was clearly different from the others - resistant to ceftazidime, cefotaxime, ceftriaxone, and non-susceptible to cefoperazone and cefepime based on disc testing (**Table 1**). MIC testing confirmed this difference for ceftazidime and cefepime, extending it into 3GC/4GCs licenced for use in cattle in the UK (**Table 2**). Relative to a non-AmpC hyper-producing control *E. coli* 17, all putative AmpC hyper-producers were non-susceptible to ceftazidime and ceftiofur (a 3GC used on several study farms during the period of sample collection) but not generally cefoperazone, cefepime or cefquinome (a 4GC used on some study farms during the period of sample collection). The MICs of the 4GCs cefepime and cefquinome were, respectively, 6 and 7 doublings higher against the isolate from Farm 1 than against the control isolate, and 5 doublings higher for each drug than against the isolate from Farm 2 (**Table 2**).

**Table 1.**
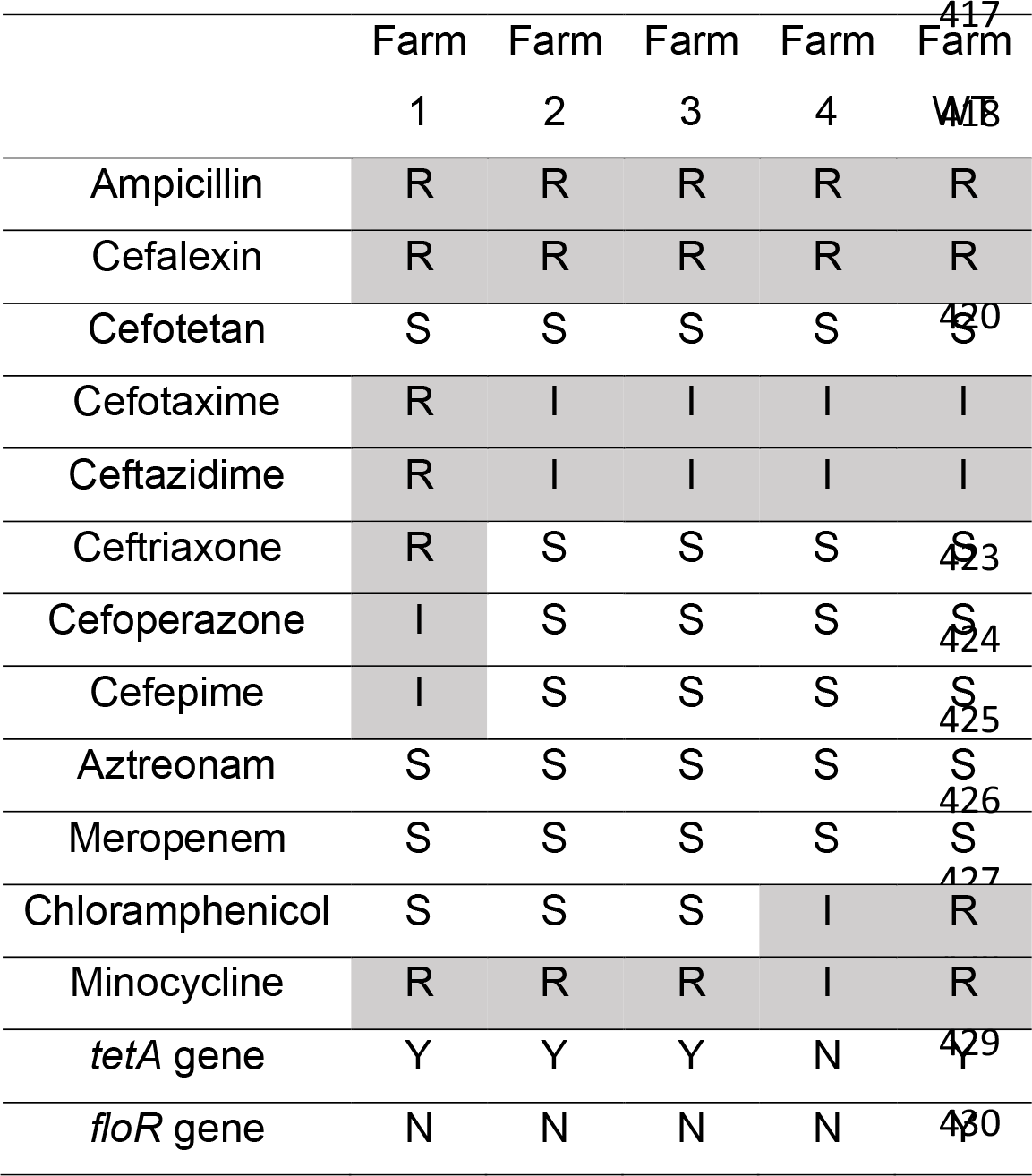
Disc susceptibility for putative AmpC hyper-producing *E. coli* isolates from dairy farms.

**Table 2.**
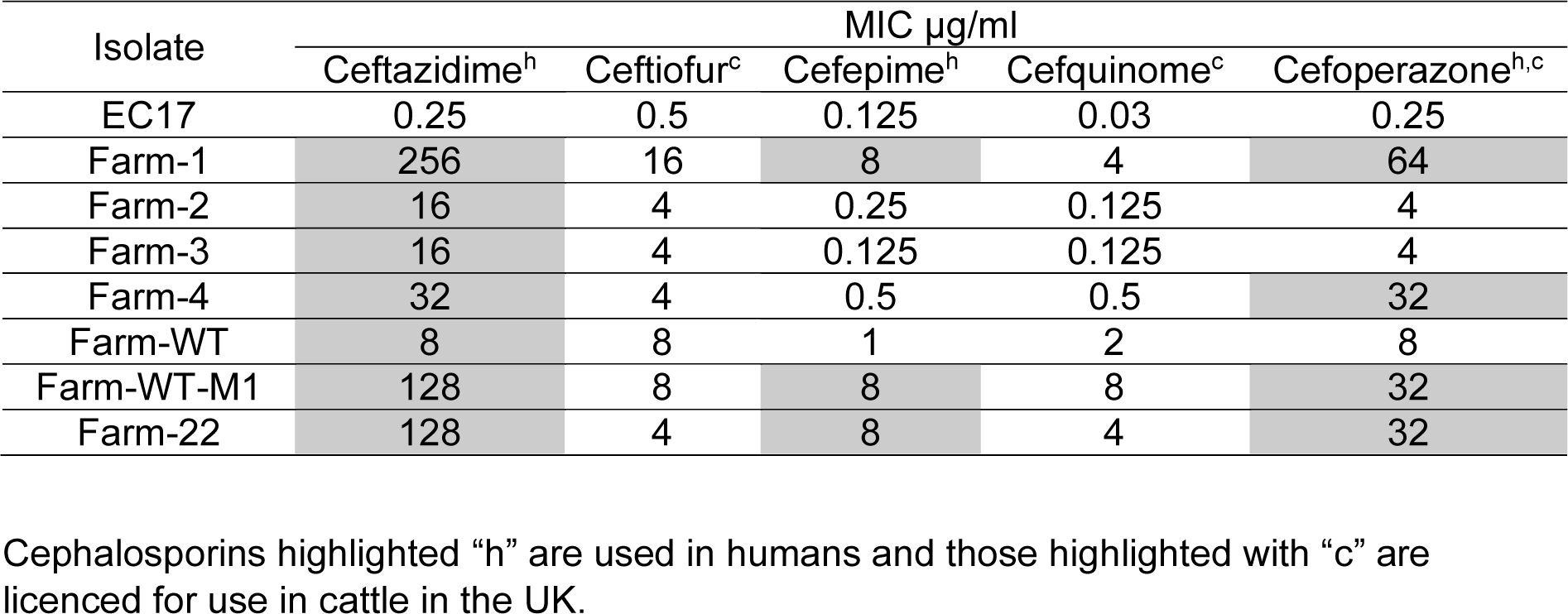
MICs of 3GC/4GCs against putative AmpC hyper-producing *E. coli* isolates from dairy farms.

| Isolate | MIC µg/ml |  |  |  |  |
| --- | --- | --- | --- | --- | --- |
|  | Ceftazidime<sup>h</sup> | Ceftiofur<sup>c</sup> | Cefepime<sup>h</sup> | Cefquinome<sup>c</sup> | Cefoperazone<sup>h,c</sup> |
| EC17 | 0.25 | 0.5 | 0.125 | 0.03 | 0.25 |
| Farm-1 | 256 | 16 | 8 | 4 | 64 |
| Farm-2 | 16 | 4 | 0.25 | 0.125 | 4 |
| Farm-3 | 16 | 4 | 0.125 | 0.125 | 4 |
| Farm-4 | 32 | 4 | 0.5 | 0.5 | 32 |
| Farm-WT | 8 | 8 | 1 | 2 | 8 |
| Farm-WT-M1 | 128 | 8 | 8 | 8 | 32 |
| Farm-22 | 128 | 4 | 8 | 4 | 32 |

Cephalosporins highlighted “h” are used in humans and those highlighted with “c” are licenced for use in cattle in the UK.

Using LC-MS/MS proteomics, AmpC hyper-production was confirmed in the isolate from Farm 1, relative to a control *E. coli* 17, but AmpC production in this isolate was not more than in the other 4 confirmed AmpC hyper-producing isolates (**Table 3**). Sequencing the *ampC* promoter region revealed that all 5 AmpC hyper-producers had the same mutations, relative to the *E. coli* 17 control (**Figure 1**), which have previously been shown to cause *ampC* hyper-expression.^1^ Proteomics showed that, unlike the other 4 AmpC hyper-producers, the cefepime-resistant isolate from Farm 1 did not produce the OmpF porin (**Table 3**), and WGS revealed a loss of function mutation in *ompF* caused by the insertion of IS*4* at nucleotide 625. OmpF porin loss did not noticeably affect envelope permeability in the Farm 1 isolate relative to the other 4 isolates or the *E. coli* 17 control (**Figure 2**). Indeed, the isolate from Farm 4 had markedly reduced permeability, reminiscent of an efflux hyper-production phenotype (constant reduced accumulation of the fluorescent dye; **Figure 2**) and yet it was not resistant to cefepime (**Table 2**). Proteomics confirmed hyper-production of AcrAB-TolC in the Farm 4 isolate and down regulation of OmpF porin (**Table 3**). This was reminiscent of a Mar phenotype and suspected loss of function mutation in *marR* was confirmed by WGS (causing a Pro57Thr change in MarR). As expected of a Mar isolate, the Farm 4 isolate was non-susceptible to minocycline and chloramphenicol, which are known AcrAB-TolC substrates. Other isolates resistant to these agents, were shown by WGS to carry specific mobile resistant genes (**Table 1**). Interestingly, the Farm 4 isolate was cefoperazone-resistant (**Table 2**). It would seem, therefore, that a combination of AmpC plus AcrAB-TolC hyper-production and/or OmpF down regulation leads to cefoperazone resistance in *E. coli*. Cefoperazone has been, albeit rarely, used as a therapy for mastitis in dairy cows in the UK.

**Table 3.** Abundance of key resistance proteins in putative AmpC hyper-producing *E. coli* from dairy farms and human urinary tract infectionstitle. Protein abundance is reported relative to the average abundance of ribosomal proteins in a cell extract and is a mean +/− standard error of the mean, (n=3). Proteins whose abundance is significantly (p<0.05) up or downregulated at least 2-fold relative to the EC17 control (see methods) are shaded in red or green, respectively.

| Accession | Description | EC17 | Farm-1 | Farm-2 | Farm-3 | Farm-4 | Farm-WT | Farm-WT-M1 | UTI-8 | UTI-9 |
| --- | --- | --- | --- | --- | --- | --- | --- | --- | --- | --- |
| P02931 | OmpF | 0.69<br>±0.36 | 0.02*<br>±0.03 | 0.99<br>±0.36 | 1.03<br>±0.34 | 0.12*<br>±0.08 | 1.54<br>±1.34 | 0.81<br>±0.24 | 0.86±<br>0.18 | 0.43±<br>0.31 |
| P00811 | AmpC | ND | 0.79*<br>±0.19 | 0.86*<br>±0.20 | 0.89*<br>±0.16 | 0.96*<br>±0.20 | 1.13*<br>±0.77 | 0.76*<br>±0.24 | 2.13±<br>0.37 | 1.35±<br>0.34 |
| P0AE06 | AcrA | 0.10<br>±0.04 | 0.13<br>±0.05 | 0.18<br>±0.15 | 0.11<br>±0.03 | 0.20*<br>±0.01 | 0.13<br>±0.07 | 0.12<br>±0.02 | 0.14±<br>0.02 | 0.16±<br>0.03 |
| P31224 | AcrB | 0.07<br>±0.01 | 0.07<br>±0.06 | 0.14*±<br>0.03 | 0.08<br>±0.08 | 0.11*<br>±0.02 | 0.04<br>±0.01 | 0.05<br>±0.01 | 0.07±<br>0.02 | 0.07±<br>0.02 |
| P02930 | TolC | 0.12<br>±0.06 | 0.08<br>±0.07 | 0.13<br>±0.02 | 0.12<br>±0.02 | 0.39*<br>±0.09 | 0.19<br>±0.10 | 0.19<br>±0.05 | 0.16<br>±0.03 | 0.10<br>±0.04 |

**Figure 1.**
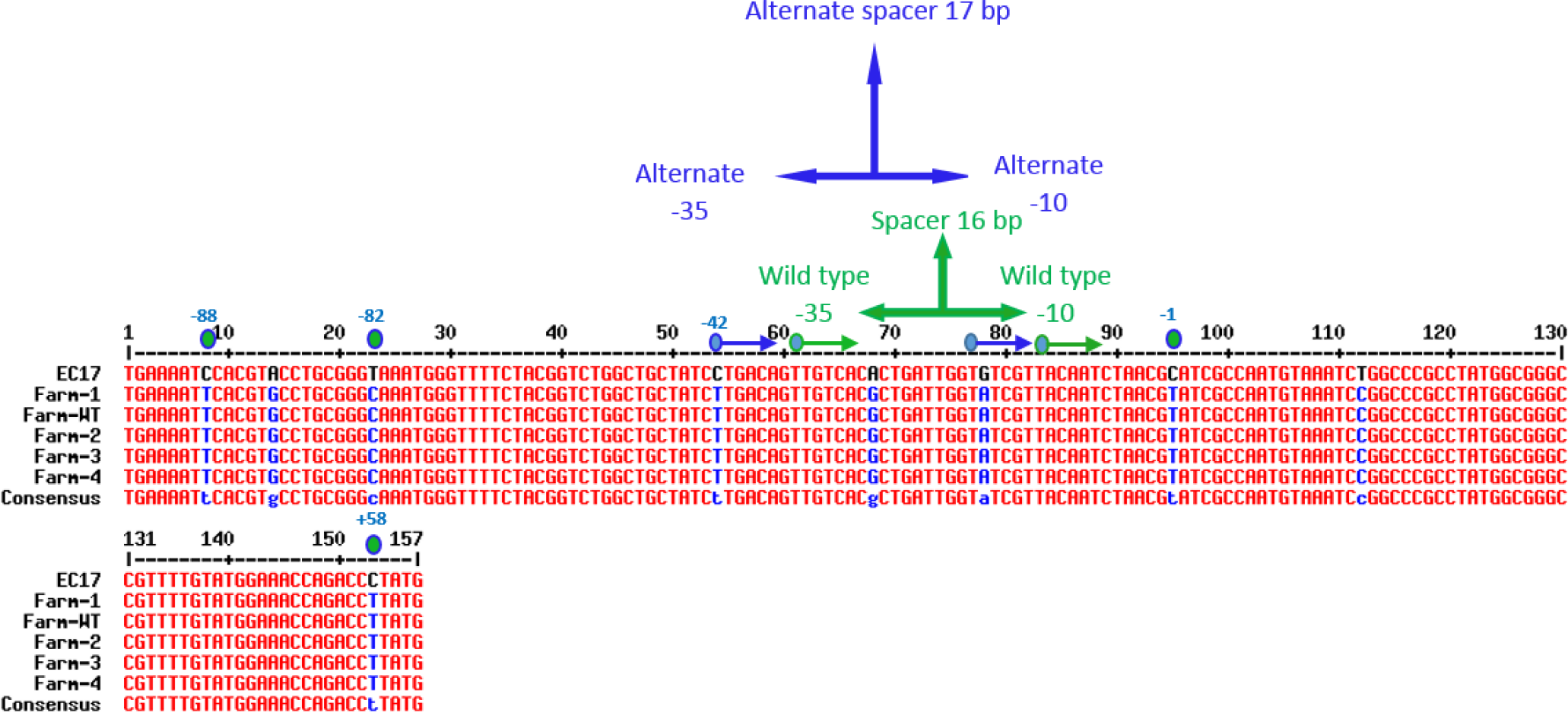
Promoter/attenuator sequences for *ampC* from 5 farm *E. coli* AmpC hyper-producing isolates in comparison with a wild-type *E. coli* EC17. Modified residues relative to the transcriptional start site are noted and the novel promoter created is annotated.

**Figure 2.**
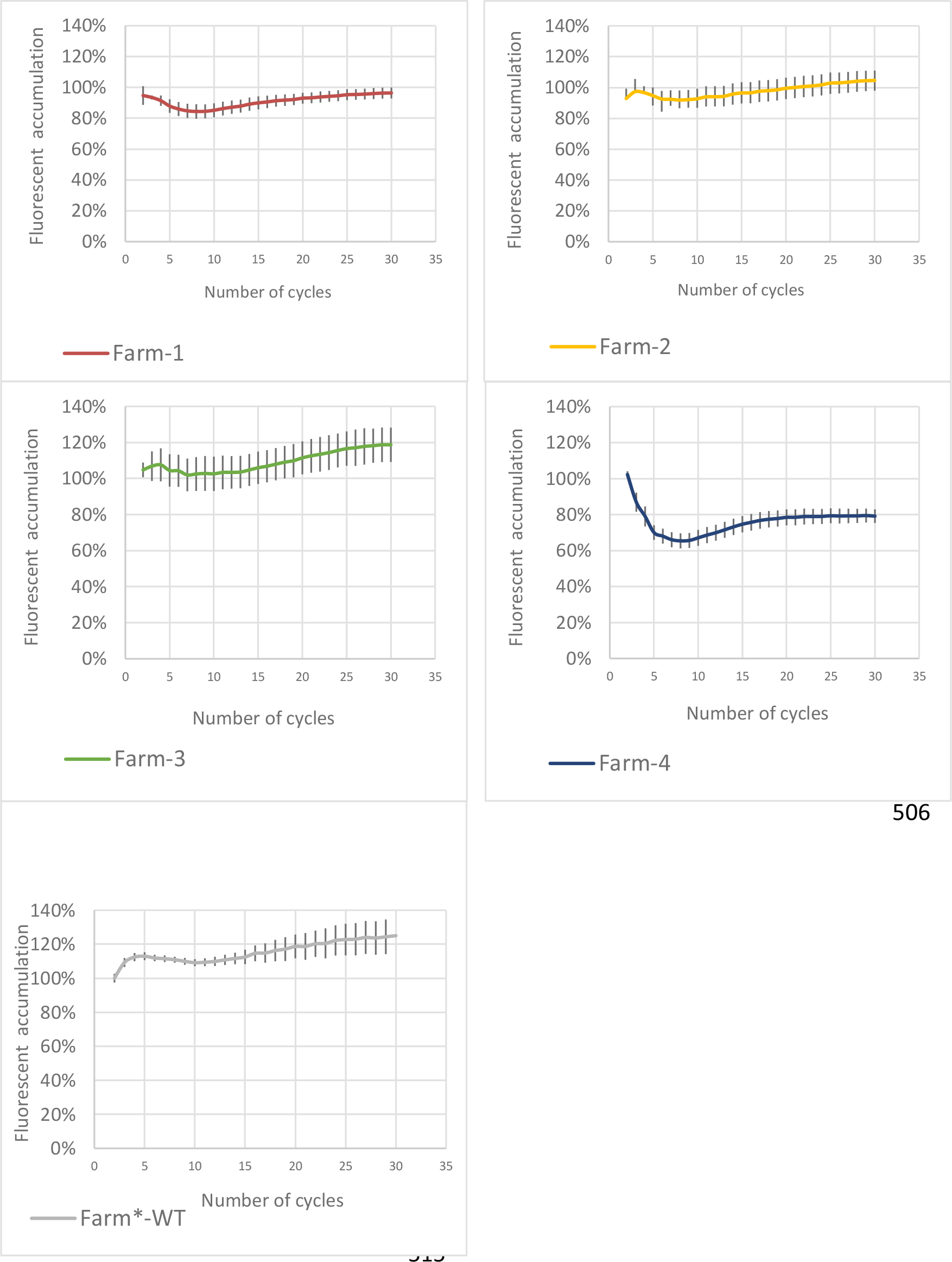
Envelope permeability of AmpC hyper-producing *E. coli* determined using fluorescent dye accumulation assays. In each case, fluorescence of an AmpC hyper-producing isolates (Farm-1, -2, etc.) incubated with the dye is presented relative to that in the control *E. coli* strain EC17 after each cycle. Each line shows mean data for 3 biological replicates with 8 technical replicates in each. Error bars define the standard error of the mean.

### First identification of expanded-spectrum AmpC variants in *E. coli* from UK dairy farms and phylogenetic analysis of AmpC hyper-producers showing recent transmission between farms

Having ruled out additional AmpC hyper-production as the cause of 4GC and cefoperazone resistance in the isolate from Farm 1, we next looked at the *ampC* gene sequence. There were several sequence nucleotide polymorphisms from one *ampC* gene to the next amongst our 5 representative isolates, but only one in the Farm 1 isolate stands out: causing a His312Pro change (His296Pro when considering the mature AmpC protein following removal of the signal peptide), a mutation previously shown to enhance the spectrum of AmpC hydrolytic activity.^31^

WGS showed that isolate Farm-WT had an identical *ampC* open reading frame and promoter sequence to that carried by the isolate from Farm 1, but without the single mutation predicted to cause expanded-spectrum AmpC activity. We therefore selected a mutant (Farm-WT-M1) using ceftazidime at its agar dilution MIC (8 mg/L) using Muller Hinton Agar. The mutant did not have altered production of key resistance proteins relative to its parent, Farm-WT (**Table 3**). Sequencing of the *ampC* gene from Farm-WT-M1 revealed an identical His296Pro mutation to that seen in the isolate from Farm 1, and the mutant had the same expanded-spectrum antibiogram as the isolate from Farm 1 (**Table 2**). Since Farm-WT-M1, like its parent, has wild type *ompF*, this confirmed that the insertional inactivation of *ompF* seen in the isolate from Farm 1 had little impact on the MICs of expanded-spectrum cephalosporins in the presence of an expanded-spectrum AmpC variant (**Table 2**).

We next performed WGS analysis of putative AmpC hyper-producers identified in our molecular epidemiology survey ^11^ from 25 farms across the core portion of our study area;^10^ this area also included the locations of 146 GP practices involved in a parallel survey of human urinary *E. coli*.^3^ All 25 representative isolates had the same *ampC* promoter mutation reported above (**Figure 1**). **Table 4** shows the spread of *E. coli* STs. Similar to a reported cattle study in France,^8^ ST88 was dominant (10/25 isolates). Based on analysis of *ampC* sequence, only one other isolate (from Farm 22) was found to carry a known expanded-spectrum AmpC variant, in this case with the same His296Pro mutation as seen in the isolate from Farm 1. This isolate had the same expanded spectrum antibiogram as that from Farm 1 (**Table 2**). These 2 isolates, from farms 40 km apart, were both ST641 and only 64 SNPs apart in the core genome, based on phylogenetic analysis (**Figure 3**). This can be compared with SNP distances of 1-13 SNPs across 6 sequenced isolates collected from Farm 1 over a 12-month period. Interestingly, the *ompF* porin gene was intact in the isolate from Farm 22 so *ompF* disruption must have occurred following separation of the isolates. Measurement of MICs against the isolates provided further evidence that loss of *ompF* was not important for 3GC/4GC resistance conferred by the expanded-spectrum AmpC in the isolate from Farm 1 (**Table 2**). Interestingly, another ST641 isolate, from Farm 7 (which is 7 km from Farm 1), had 1520 SNPs different from the isolate from Farm 1 (**Figure 3**) and did not have the expanded-spectrum AmpC mutation or an *ompF* mutation; this isolate shared these properties with the isolate from Farm 14, which was only 35 SNPs (**Figure 3**) but 45 km away from Farm 7.

**Table 4.** Sequence types of representative AmpC hyper-producing isolates from 25 dairy farms.

| **Isolate** | **ST** | **Phylogroup** |
| --- | --- | --- |
| Farm-1 | 641 | B1 |
| Farm-2 | 88 | C |
| Farm-3 | 88 | C |
| Farm-4 | 388 | B1 |
| Farm-5 | 88 | C |
| Farm-6 | 75 | B1 |
| Farm-7 | 641 | B1 |
| Farm-8 | 23 | C |
| Farm-9 | 162 | B1 |
| Farm-10 | 88 | C |
| Farm-11 | 2522 | B1 |
| Farm-12 | 88 | C |
| Farm-13 | 278 | B1 |
| Farm-14 | 641 | B1 |
| Farm-15 | 88 | C |
| Farm-16 | 278 | B1 |
| Farm-17 | 661 | B1 |
| Farm-18 | 88 | C |
| Farm-19 | 88 | C |
| Farm-20 | 278 | B1 |
| Farm-21 | 345 | B1 |
| Farm-22 | 641 | B1 |
| Farm-23 | 88 | C |
| Farm-24 | 278 | B1 |
| Farm-25 | 88 | C |

**Figure 3.**
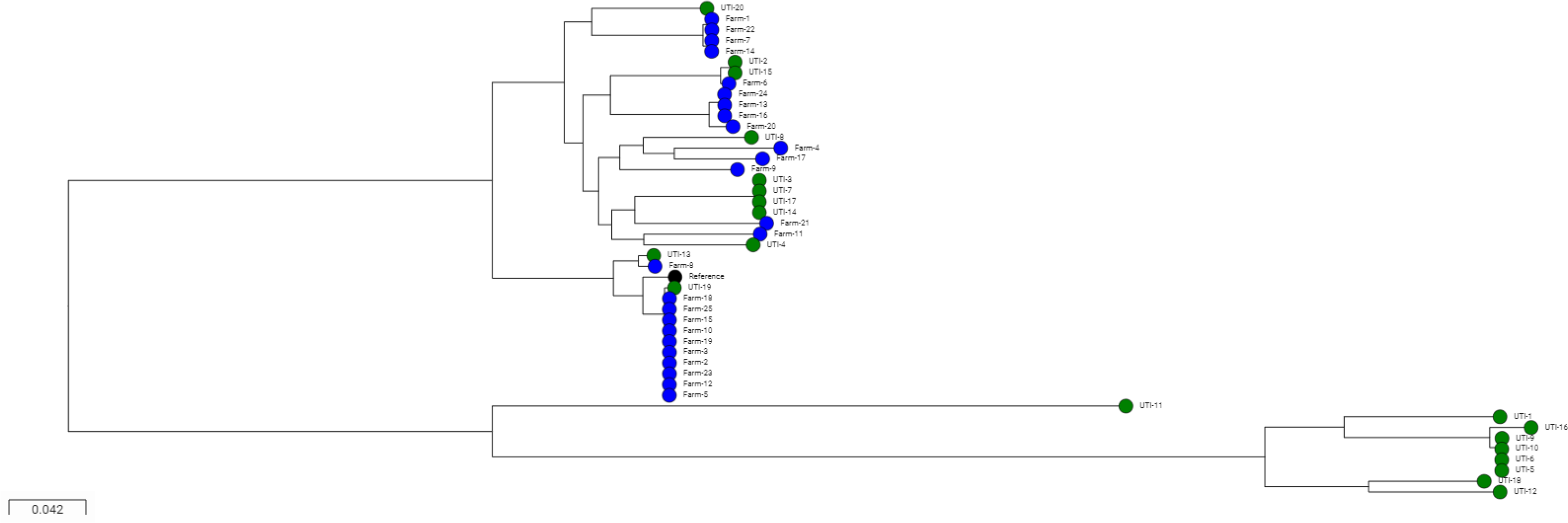
Phylogenetic tree of farm and human urinary AmpC hyper-producing *E. coli*. The phylogenetic tree was illustrated using the Microreact program using a maximum likelihood tree generated from core genome alignments as described in Materials and Methods. Isolates are coloured green (human urinary) and blue (farm). The ST88 finished reference genome used to generate the alignments is noted in black.

### Risk factor analysis

The data presented above, when considered in conjunction with that in our recent survey,^11^ show that 46.2% of CTX-R *E. coli* from dairy cattle across the 53 farms enrolled in our study were AmpC hyper-producers. This compares with 52.9% that were CTX-M producers, the remainder being plasmid AmpC producers.^11^ Accordingly, attempts to reduce the prevalence of 3GC resistance on dairy farms must address the specific factors that are driving the accumulation of AmpC hyper-producers. In order to identify factors associated with an increased risk of finding CTX-R, AmpC hyper-producing *E. coli* in a sample from farms in our study, we performed risk factor analyses. Three farm-level fixed effects and 2 sample-level fixed effects were identified as important (**Table 5**). As seen with our risk factor analysis for *bla*_CTX-M_-positive CTX-R *E. coli* on the same farms,^10^ samples collected from the environment of young calves were much more likely to be positive for AmpC hyper-producing *E. coli* (p<0.001) and samples collected from pastureland, including publicly accessible sites, were much less likely to be positive (p=0.005). We found no association between cephalosporin use – including 3GC use – and increased risk of finding AmpC hyper-producers. Interestingly, however, the total usage of amoxicillin-clavulanate was associated with a higher risk of finding AmpC hyper-producing *E. coli* on a farm (p=0.001). This association can be explained by direct selection since AmpC hyper-production confers amoxicillin-clavulanate resistance in *E. coli*.^1^ This finding is important because amoxicillin-clavulanate is not currently identified as a highest-priority critically important antimicrobial (HP-CIA) by the World Health Organisation,^32^ and, whilst great strides have been made within the UK farming industry to reduce antibiotic use,^33^ there is a particular focus on reducing HP-CIA, e.g. 3GC use. The associations identified in our risk factor analysis suggest that reducing HP-CIAs without also reducing amoxicillin-clavulanate use may not impact on the prevalence of CTX-R, AmpC hyper-producing *E. coli* on farms. Indeed, a bigger concern is that reducing 3GC use on farms may drive up amoxicillin-clavulanate use providing additional co-selective pressure for 3GC-resistant *E. coli*.

**Table 5.** Significant associations (p<0.05) with AmpC hyper-producing *E. coli* from dairy farms from the multilevel, multivariable logistic regression model

| **Risk factor** | **Odds ratio [95% confidence interval]** | **p** |
| --- | --- | --- |
| Sample taken from the environment of pre-weaned heifers | 3.93 [2.72, 5.67] | <0.001 |
| Total usage of amoxicillin-clavulanate on the farm | 1.16 [1.06, 1.27] | 0.001 |
| Routine use of vaccination against respiratory disease in calves | 2.72 [1.34, 5.56] | 0.005 |
| Samples taken from pastureland | 0.32 [0.14, 0.72] | 0.005 |
| Calving all-year-round as opposed to in seasonal blocks | 3.97 [1.45, 10.86] | 0.006 |

A final observation from this analysis is that average monthly temperature, which was identified as a strong risk factor for finding *bla*_CTX-M_-positive *E. coli* in this same survey of dairy farms,^10^ was not identified as a risk factor for finding AmpC hyper-producing *E. coli*. This may be an issue of power, but the numbers of *bla*_CTX-M_ *E. coli* positive and AmpC hyper-producing *E. coli* positive samples in the survey were similar (224 vs 186). It may be hypothesised, therefore, that carriage of (i.e. because of some fitness cost) or transmission rate for the horizontally acquired *bla*_CTX-M_ is specifically affected by temperature, whereas the presence of chromosomal mutations in the *ampC* promoter leading to AmpC hyper-production is not.

### No evidence for recent human/farm transmission of AmpC hyper-producing *E. coli* isolates collected in parallel in a 50 × 50 km region

We next looked at WGS data for 20 human urinary *E. coli* presumed to hyper-produce AmpC, collected during the same timeframe from people living in the same geographical range as the 25 farms for which WGS data of AmpC hyper-producing *E. coli* had been obtained.^3^ Proteomics confirmed AmpC hyper-production in 2 representative isolates: UTI-8 and UTI-9, with almost double the amont of AmpC in UTI-8 than in UTI-9 (**Table 3**). There were 9 different *ampC* promoter types seen across the 20 AmpC hyper-producing human isolates, though 11/20 isolates carried the same promoter mutation seen in all 25 farm isolates, including UTI-8 and UTI-9, though UTI-8 also has an attenuator mutation at +37, which probablhy explain the higher level of AmpC seen in UTI-8 that UTI-9 (**Figure 4**, **Table 3**). None of the human isolates had mutations suggestive of an expanded spectrum AmpC variant, which was confirmed phenotypically using cefepime disc susceptibility testing.

**Figure 4.**
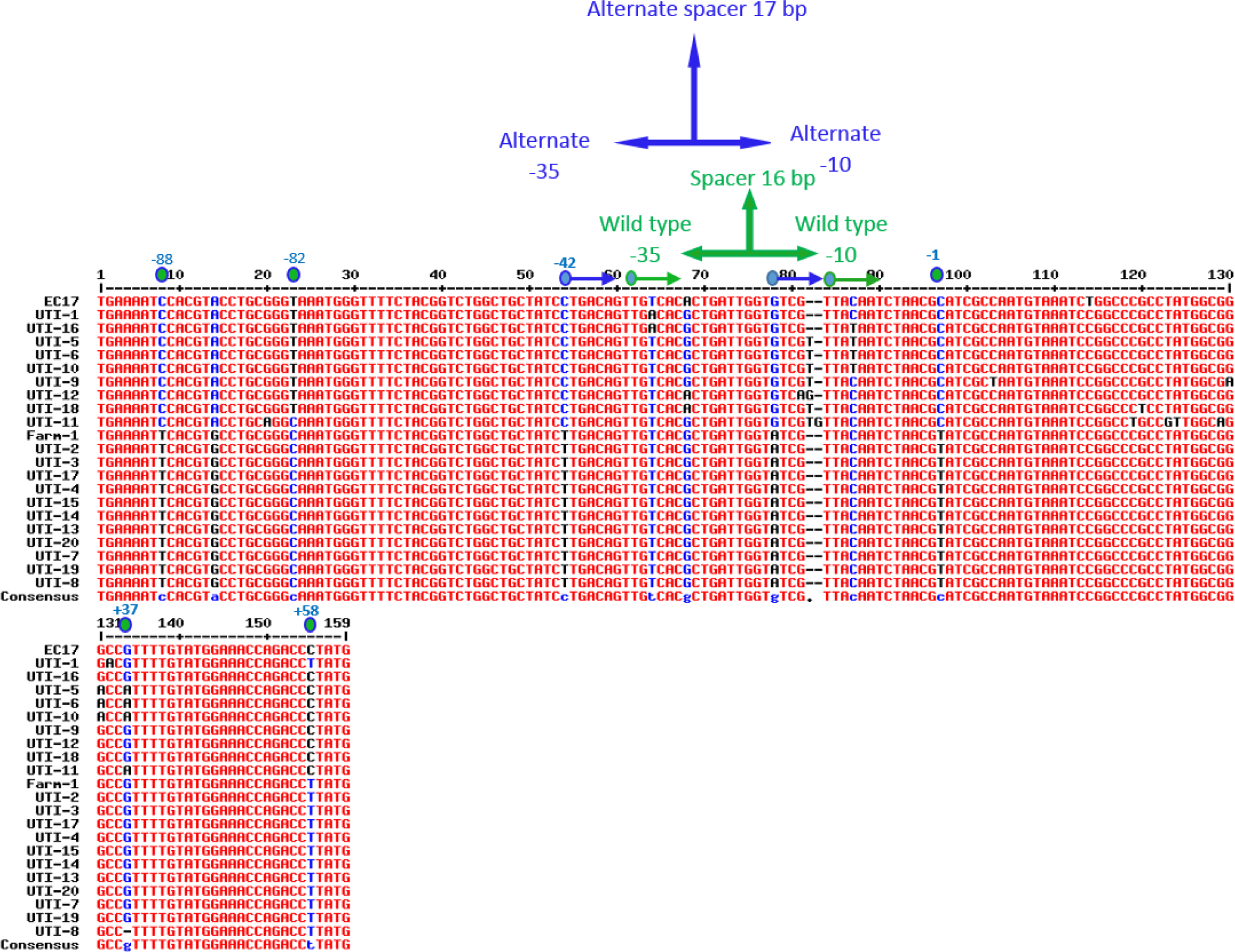
Promoter/attenuator sequences for *ampC* from 20 human urinary *E. coli* AmpC hyper-producing isolates in comparison with a wild-type *E. coli* EC17 and the isolate from Farm 1. Modified residues relative to the transcriptional start site are noted and the novel promoter created is annotated.

Our final aim was to identify if there was any evidence of sharing AmpC hyper-producing *E. coli* between humans and cattle, since dominance of ST88 has previously been reported in humans in Northern Europe ^9^ and since we found an over-representation of ST88 on our farms (**Table 4**). A phylogenetic tree drawn based on core genome comparison showed that the cattle and human isolates were intermixed only to a small extent, with only one human ST88 isolate found (**Figure 3**). Importantly, all 10 ST88 cattle isolates were 15 or fewer SNPs apart, suggesting very recent farm-to-farm transmission; the human ST88 isolate (UTI-19) was, at its closest distance, 1279 SNPs different from the cattle isolates. The 2 other examples where isolates from the same ST were found in farm and human samples gave the same story (**Figure 3**): for ST75, the 2 human isolates (UTI-2 and UTI-15) were 60 SNPs apart, but the cattle isolate (Farm-6) was 1972 SNPs different at best. For ST23, the human and cattle isolates (UTI-13 and Farm-8, respectively) were 2754 SNPs different. Otherwise, there was no ST sharing, and all cattle isolates fell into phylogroups B1 and C (**Table 4**), with 8/20 human isolates falling into the highly pathogenic phylogroup B2, including a cluster of ST73 isolates of which 3 were only 2 SNPs apart.

### Conclusions

AmpC hyper-production is a remarkably common mechanism of 3GC resistance in *E. coli* from dairy farms in our study - similar to a national survey in The Netherlands.^12^ We have shown an association between amoxicillin-clavulanate use and the risk of finding AmpC hyper-producers on dairy farms and would caution against a blanket switch from 3/4GCs to amoxicillin-clavulanate in response to justifiable action to reduce HP-CIA use. However, our comparison between AmpC hyper-producing farm and human urinary *E. coli* in the same region provided no evidence of local sharing of AmpC hyper-producers between farms and the local human population. Accordingly, whilst reducing the on-farm prevalence of AmpC hyper-producing *E. coli* should be an important aim, the primary reason for achieving this would be to reduce the likelihood of difficult to treat infections in cattle rather than because of any direct zoonotic threat.

## Acknowledgements

Genome sequencing was provided by MicrobesNG (http://www.microbesng.uk), which is supported by the BBSRC (grant number BB/L024209/1).

## Funding

This work was funded by grant NE/N01961X/1 to M.B.A., K.M.T., D.C.B. and K.K.R. from the Antimicrobial Resistance Cross Council Initiative supported by the seven United Kingdom research councils. M.A. was in receipt of a postgraduate scholarship from the Saudi Cultural Bureau.

## Transparency declaration

The authors declare no conflict of interests. Farming and veterinary businesses who contributed data and permitted access for sample collection were not involved in the design of this study or in data analysis and were not involved in drafting the manuscript for publication.

